# Arsenic Impairs Critical Cellular Events of Human Cardiac Development Across Differentiation and Metabolic Maturation

**DOI:** 10.64898/2026.10.11.758344

**Authors:** Bailey Renee Skeen, Cate Duncan, Fanrong Yao, Brian Shewchuk, Karen Mruk, Erik J. Tokar, Xian Wu

## Abstract

Arsenic (As) is a widespread environmental contaminant, with exposure occurring primarily through contaminated food and water or cigarette smoke. Maternal exposure to As represents a significant risk factor for congenital heart disease in offspring. However, the mechanisms underlying As-induced developmental cardiotoxicity remain poorly defined. Pluripotent stem cells provide an excellent platform for studying mechanisms of developmental toxicity. Embryoid bodies (EBs) derived from stem cells, can recapitulate many key events of early embryogenesis including germ layers specification. Human cardiomyocytes are derived from mesodermal progenitors. NKX2- 5 governs cardiac specification and morphogenesis, and its mutation produces atrial and ventricular defects in humans. Using an NKX2-5 reporter stem cell line, we modeled mesoderm differentiation in EBs and cardiac induction in a 2D cardiomyocyte differentiation system. In an 8-day differentiation protocol, As (0.05 μM; 3.75 ppb) significantly suppressed cardiomyocyte differentiation and cardiac induction. However, As increased mesodermal markers (HAND1, SNAI2, HOPX) gene expression. In contrast, cardiac-specific genes, including NKX2-5, GATA4, Troponin T, and α-Actinin, were markedly downregulated in the presence of As. As also reduced expression of genes related to cardiac induction, fibroblast function, and vascular markers in the cardiac organoid model. As impaired cardiomyocyte maturation related gene expressions at concentrations without affecting cardiomyocyte proliferation. In a zebrafish model of early heart development, As increased heart rate and disrupted ion channel gene expression. In conclusion, As inhibits ESC differentiation into cardiomyocytes and disrupts cardiomyocyte maturation. Combining stem cell–derived models with zebrafish exposure, this study illuminates adverse outcome pathways for As-induced congenital heart defect.

## Introduction

Cardiac development is tightly programmed from heart tube formation to looped, chambered, and electrically coupled organ [1]. It is driven by conserved signaling and rapid shifts in cellular metabolism and biomechanics. Brief disruptions during these windows can lead to heart developmental defects like congenital heart diseases [2]. Exposure during heart development that disrupts the developmental stages leading to either observable congenital heart defects or more subtle epigenetically encoded immaturity leading to increased cardiovascular risk across the lifetime. Integrating human stem cells-derived 2D cardiomyocytes and 3D cardiac organoids with multi- parameter assays (e.g., contractility, electrophysiology, metabolic flux, and transcriptomics) will provide a comprehensive and human-relevant approach to assess how environmental exposures perturb cardiac development and long-term cardiovascular health [3].

Human stem cell-derived cardiomyocytes and 3D organoids now enable time-resolved, mechanistic studies with functional and multi-omic readouts, providing the basis for biomarkers and interventions that protect the developing heart. Embryonic stem cells (ESC) or induced pluripotent stem cells (iPSC) derived cardiomyocytes are utilized as in vitro models for cardiac toxicology studies [4]. Conventional 2D ESC models via embryoid bodies (EBs) in suspension medium or a monolayer of ESC or iPSC cells in matrigel are commonly employed for cardiac differentiation in the contexts of cardiovascular disease models, drug screening, development, and cell therapy applications [5]. In a related investigation, 81 drugs were examined using commercially available iPSCs over 72 hours, utilizing 0.03-100 μM doses and a metabolic biomarker-based assay to anticipate variations in metabolism and viability of iPSC-CMs [6]. Human iPSC engineered cardiac tissues demonstrated optical simulation by augmenting the heart rate of human iPSC-CMs to enhance maturation and improve electrophysical properties of the cardiac tissue [7]. The NKX2-5 homeobox transcription factor, essential for almost all stages of cardiac development and adulthood, when mutated, can trigger various congenital heart abnormalities [8]. To monitor heart development in real-time and characterize the dynamic gene-expression changes underlying cardiomyocyte differentiation, cardiomyocyte reporter hESC lines have been developed using CRISPR-Cas technology. These types of models serve facilitating a comprehensive understanding of arsenic’s impact on fetal cardiac development.

Nonetheless, 2D cell cultures have limitations such as changed morphology due to flattening, increased proliferation rate, decreased ability to contract or form cell-to cell junctions and interactions, and lack of environmental factors, which can lead to immature cardiomyocytes [9]. Despite their affordability, easy maintenance, and similarities in cardiac function, gene and protein expression, rodents are not ideal models for translational studies in human clinical trials due to their different heart rates, blood pressures, contractility, and ventricular cardiomyocyte expressions compared to humans [10]. Thus, 3D cardiac organoid models, with their ability to mimic early in vivo heart development and facilitate cell-to-cell and cell-to-matrix interactions, have emerged as superior models for studying developmental cardiotoxicity [11, 12].

Arsenic (As), an environmental metalloid, has been identified as a significant contributor to congenital heart defects in newborns across the globe [13, 14]. Exposure to arsenic during fetal development may arise from multiple sources, including drinking water, food consumption, industrial emissions, and maternal smoking, potentially causing long-term damage to the cardiovascular system. In an epidemiological study, it was found that arsenic can cross the placental barrier, reaching a median concentration of 34ug/kg in the unborn child, and further persisting after birth as dimethylarsinic acid in women exposed to arsenic during pregnancy [15]. A study on rodents involving 21 days of intravenous arsenic trioxide injections and treatment of H9c2 rat cardiac cells with varying concentrations of arsenic trioxide unveiled both in vivo and in vitro upregulation of mitogen activated protein kinase, leading to cardiac cell death [16]. With an increasing trend of cardiovascular defects in newborns, studies in cardio toxicology are being conducted to understand the impact of environmental metals such as arsenic on the body. Evidence shows that arsenite exposure escalates reactive oxygen species, activating inflammatory cytokines that cause atherosclerosis, and impairs nitric oxide via the endothelial nitric oxide synthase, potentially leading to cardiovascular anomalies due to vascular endothelial dysfunction. However, the mechanisms by which arsenic regulates cellular metabolism and cell differentiation during heart development remain incompletely understood.

## Materials and Methods

### Cell Maintenance and Differentiation

NKX2-5^eGFP/w^ hESCs were kindly gifted from Dr. E.G. Stanely lab and cultured on Matrigel® (354230; Corning, NY) coated plates in mTeSR™ Plus medium (100-0276; STEMCELL Technologies Inc, Canada) until 80-90% confluent before cell passaging and differentiation. Cardiomyocyte differentiation was performed in 96-well plates (353219; Corning, NY) using STEMdiff™ Cardiomyocyte Differentiation Kit (05010; STEMCELL Technologies Inc, Canada) as described in the protocol. Typically, on day 0, the NKX2-5eGFP/w hESCs were seeded on Matrigel® coated plates at density of 3 x 10^4^ cells per well in 200 μl of STEMdiff™ Cardiomyocyte Differentiation Medium A and incubated at 37°C for 2 days. On day 2, medium was gently removed, 200 μl of STEMdiff™ Cardiomyocyte Differentiation Medium B was added and plates were incubated at 37°C for 2 days. On day 4 and again on day 6, medium was gently removed, 200 μl of STEMdiff™ Cardiomyocyte Differentiation Medium C was added and plates incubated at 37°C for 2 days. On day 8, medium was removed, cells were washed with PBS before cell collecting and analysis.

Formation of 3D -EBs-and differentiation to germ layers was performed in 384-well plates using Essential 6™ Medium (A1516401; ThermoFisher Scientific, MA) with a cell density of 1000 cells/well for 8 days. A single EB was formed in each well. Sodium arsenite ( 0.001 – 100 µM, S7400, MilliporeSigma, Burlington, MA) was added together with each medium change every 48 hours from day 0 to day 8.

### Cardiac organoid formation

Accutase cell detachment solution (07920; STEMCELL Technologies Inc, Canada) was used to dissociate NKX2-5eGFP/w hESCs for organoid formation. After dissociation, cells were resuspended in Essential 8 Flex medium containing 10 μM ROCK inhibitor Y- 27632 (S6390; Selleckchem, PA). Cells were then counted using a cell counter (1450102; Biorad, CA) and seeded at 1 x 104 cells/well in 100 μL of medium in round bottom ultra-low attachment 96-well plates (4520; Corning, ME) on day −2. The plate was then centrifuged at 1000 rpm for 5 minutes and placed in an incubator at 37°C, 5% CO2. After 24 hours (day −1), 50 μl of medium was carefully removed from each well and 200 μl of fresh Essential 8 Flex medium was added for a final volume of 250 μl/well. The plate was returned to the incubator for 24 hours. On day 0, 166μl of media was removed from each well and 166 μl of RPMI 1640/B-27, minus insulin (A369520; ThermoFisher Scientific, MA) containing CHIR99021 (S1263; Selleck) was added at a final concentration of 4 μM/well along with BMP4 at 0.36 pM (1.25ng/ml; 314-BP-010; R&D, MN) and Activin A at 0.08 pM (1ng/ml; 338-AC-010; R&D, MN) for 24 hours. On day 1, 166 μl of medium was removed and replaced with fresh RPMI1640/B-27, minus insulin. On day 2 and again on day 4, RPMI/B-27, minus insulin containing IWR-1 (S7086; Selleckchem, PA) was added for a final concentration of 5 μM IWR-1 and the cells incubated for 48 hours. On day 6, medium was changed to RPMI1640/B-27 (17504044; ThermoFisher Scientific, MA). On day 7, a second 4 μM CHIR99021 exposure was conducted for 1 hour in RPMI1640/B-27. Subsequently, medium was changed every 48 hours for organoids maintenance. Sodium arsenite solution (0.001 – 10 mM) was added every 48 hours during differentiation.

### Image Acquisition and Analysis

Cardiomyocytes derived from NKX2-5^eGFP/w^ hESCs at different days were allowed to adhere in 96-well flat clear bottom black walled polystyrene tissue-culture treated microplates (353219; Corning, ME). The plates were transferred to an IncuCyte ZOOM® platform which was housed inside an incubator at 37 °C/5% CO2 for the duration of the assay. Four images per well from two technical replicates were taken using a 10× objective and then analyzed using the IncuCyte™ Basic Software. Green channel acquisition time was 450 ms. In phase contrast, cell segmentation was achieved by applying a mask to exclude cells from background. An area filter was applied to exclude objects smaller than 50 μm^2^. Green channel background noise was subtracted with the adaptive method of background non-uniformity correction with a threshold adjustment at 0.23 green corrected units. Fluorescence signal was quantified applying a mask. Edge split was used to split closely spaced objects to obtain a more accurate quantification of fluorescent objects.

Wild-type zebrafish embryos were maintained in a 6-well plate that was kept in an incubator maintained at 28 °C. The mounted zebrafish were transferred to a Leica M165 FC microscope imaging platform. The software used to view the zebrafish was the Leica Application Suite X (LASX) version 3.8.2.27713. Ten single fish images were taken per treatment group at a 3x objective, then analyzed using Fiji ImageJ and Microsoft Excel. Images were taken under Red, Green, and Blue (RGB) brightfield channels. Ten heartbeat recordings were taken per treatment group, under RGB brightfield channels using an 8X objective for 60 s. Videos were analyzed using Fiji Image-J and Microsoft Excel.

### Zebrafish Husbandry

Adult zebrafish [wild-type AB] 3–18 months of age were maintained at 28.5°C in a recirculating system on a 14-h light/10-h dark cycle and fed daily with brine shrimp and Zeigler’s adult zebrafish diet. Embryos were obtained through natural matings. Embryos were cultured at 28°C– 30°C in E3 medium containing 5 mM NaCl, 0.17 mM KCl, 0.33 mM CaCl_2_, and 0.33 mM MgSO_4_. All chemicals were purchased from Sigma Aldrich (St. Louis, MO). All animal procedures were approved by the IACUC committee at East Carolina University.

### Arsenic Exposure of Zebrafish

Embryos were staged as described previously [17] and dosed at 24 hours post fertilization (hpf)). The embryos were dosed with 0.1 µM of As. There were three control wells and three treatment wells, each with 20 embryos. Once dosed, embryos were placed in the incubator overnight. Embryos were exposed for a total of 24 and 48 hours.

### Heart Rate Quantification in Zebrafish

To assess cardiac function, the fish were dechorionated under a microscope by carefully removing the chorion with tweezers 24 hours after exposure. Live embryos were mounted in 1% low melting point agarose on a glass depression slide. Heartbeat was recorded on a Leica M165FC stereoscope with a 1x M-series Planapo objective equipped with a Leica K7 camera. The pre-installed LasX software was used to capture thirty-second uncompressed videos. After imaging, embryos were freed from agarose, placed back in their respective conditions, and stored in the incubator. Embryos were imaged again after 48 hours of exposure. At the end of recording, embryos were released from the agarose, flash frozen in RNase free tubes, and stored at −80°C prior to RNA extraction.

Files were opened in ImageJ/FIJI software in grayscale mode without a virtual stack [18]. A circular region of interest (ROI) was drawn on the video file on the edge of the cardiac ventricle of the zebrafish larvae. Using ImageJ’s “plot-z-axis profile,” a time profile was generated via the pixel intensity change of the ROI. Peak detection and heartbeat counting was performed in MS Excel as previously described [19]. For each video, two ROIs were sampled and averaged to mediate random errors.

### Quantitative Polymerase Chain Reaction (qPCR)

RNA extraction was carried out with the RNeasy kit (74004; Qiagen, Manchester, UK) and RNA quality was verified with a ND-100 spectrophotometer (Nano Drop Technologies, DE, USA). Complementary DNA (cDNA) was reverse transcribed from 400 to 600 ng RNA using the QuantiTect Reverse Transcription kit (205311; Qiagen, Manchester, UK) according to the manufacturer’s instructions. Twenty to thirty nanograms of cDNA were used as a template in qPCR experiments using specific primers (500 nM) and Fast SYBR™ Green Master Mix as the detection chemistry (43- 856-12; ThermoFisher Scientific, MA). The thermal profile included one phase at 95 °C for 5 min, second phase of 40 cycles at 95 °C for 20 s, 60 °C for 20 s and 72 °C for 20 s, and final phase of 72 °C for 5 min. Melt curve analysis was run after every experiment. The experiments were carried out with a QuantStudio 7 Flex platform (Applied Biosystems, MA, USA) and analyzed with the QuantStudio ^TM^ software. Cycle threshold (Ct) values were determined, and relative mRNA contents were inferred from normalization of the gene of interest expression to that of the housekeeping gene (Delta Ct). Relative expression results were plotted as (2^−ΔΔCt). Zebrafish RNA extraction: RNA was extracted from pools of 25 embryos using Trizol (Ambion #15596026) and phenol:chloroform (Millipore #19K0856166). cDNA was generated using the SuperScript IV reverse transcriptase system (Invitrogen #18091150) using dT primers to target mRNA. cDNA samples were cleaned using ethanol precipitation. The mRNA expression was quantified using SYBR™ Green in realtime PCR. Relative mRNA levels were determined using the ΔΔC_t_ method. (ΔCt = Ct(target)-Ct(*gapdh*); ΔΔCt=Ct(dosed)- Ct(untreated)). The relative fold change was calculated using 2^−ΔΔCt^.

### Statistics

Cells: Cell characterization experiments were performed twice using independent cultures with n = 4-6 wells per condition per culture. For concentration-response experiments, data were normalized within experiments to corresponding control wells prior to statistical analysis. For each concentration response examined, experiments were repeated two to three times using independent cultures as described. In cell proliferation assay, experimental values are a composite of six technical (on same plate) and three biological (different plates) replicates. All data analyses for cell characterization were done using a one-way ANOVA followed by a Tukey’s test or two-tailed t-test with a significance threshold of p < 0.05 to determine if different time point means were significantly different from corresponding control means. All concentration-response experiments were analyzed using one-way ANOVA with a significance threshold of p < 0.05 followed by a Tukey’s test. Mean values ± standard deviations for all measurements are provided. Statistical analyses were performed using Graphpad Prism 10 (GraphPad Software, CA).

Zebrafish: For all zebrafish experiments, at least two breeding tanks, each containing 2 to 4 males and 2 to 4 females from separate stocks, were set up to generate embryos. Embryos from each tank were randomly distributed across tested conditions. Unfertilized and developmentally abnormal embryos were removed before experiments. Samples sizes were calculated for a 30% effect with 80% power. For all graphs, values for individual fish are plotted, and data is presented as the mean ± standard deviation (SD). For statistical testing, each distribution was assessed using the Shapiro-Wilk test. Significant differences were determined using either an unpaired student’s t-test or non-parametric Mann-Whitney t-test.

## Results

### Effect of sodium arsenite on embryoid body and mesoderm formation

The embryoid bodies (EBs) formation mimics the early germ layer formation, including the mesoderm which is required for cardiac induction. Arsenic inhibited EBs formation with a Lethal Concentration 50% (LC50) at 1.6 µM and LC10 at 0.5 µM calculated from the dose response curve after 7 days of exposure. Sodium arsenite exposure during EBs formation led to gene alteration related to mesoderm formation. *ABCA4, ESM1, HAND1, HOPX, IL6ST, PDGFRA, SNAI2, TM4SF1* were all upregulated in the scorecard assay while RGS4 was downregulated (Fig 1A). To confirm the effect, in the 2D cardiomyocyte differentiation model, sodium arsenite exposure during the first 2 days mesoderm induction leads to increasing *HAND1, HOPX* and *SNAI2* mRNA expression (Fig 1B).

**Figure 1.**
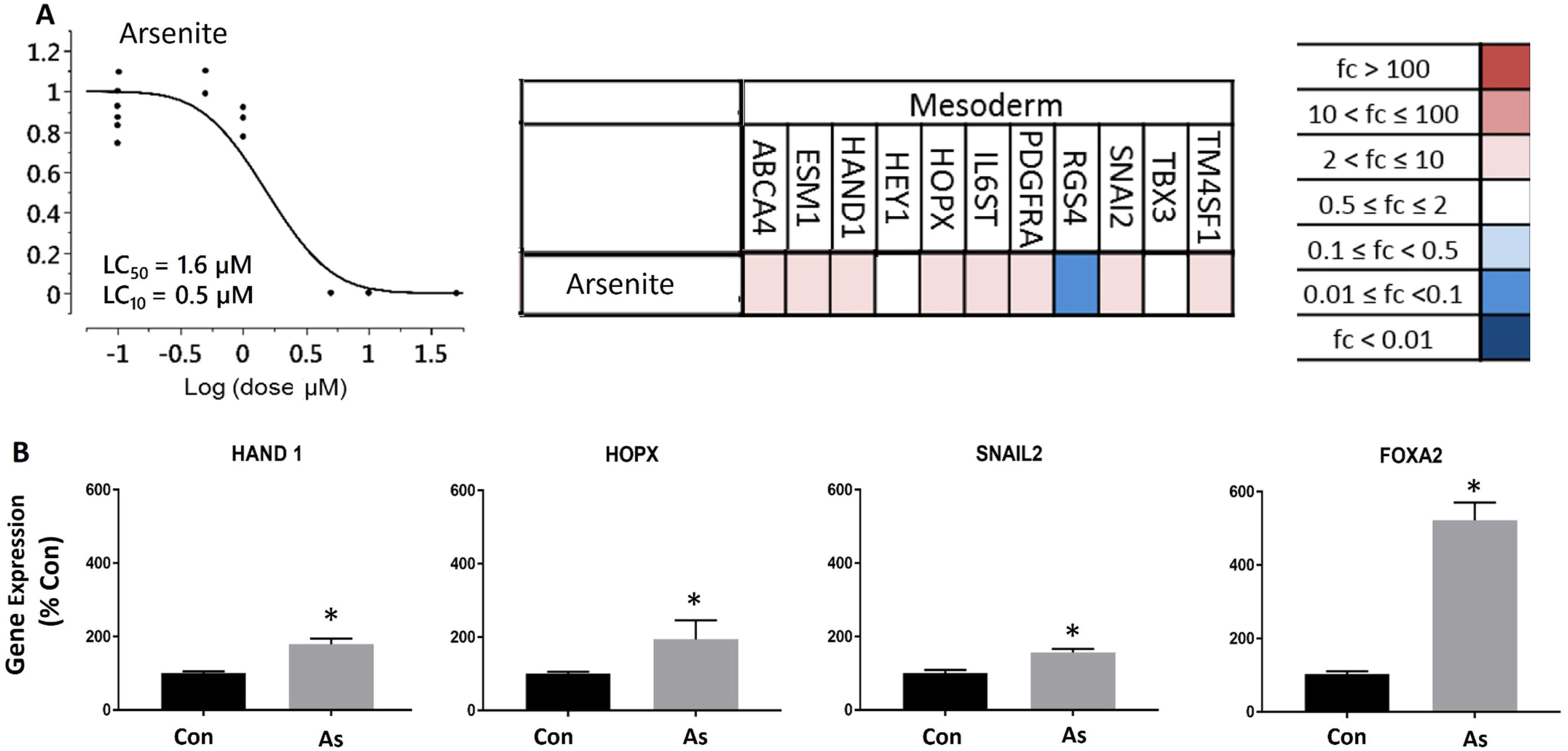
Viability and pluripotency capability of differentiating EBs exposed to As. (A) Viability was assessed using the CellTiter-Glo ATP assay (Promega) after a 96h exposure. LC50 and LC10 values were calculated using Probit analysis in JMP (version 13.0; JMP Statistical Discovery LLC). (n=8–10 per group). EB differentiation was assessed using the TaqMan hPSC Scorecard Panel following 7 d of exposure to equipotent (LC10) concen-trations. (n= 1 assay) (B) hESC 2D differentiation toward mesoderm lineage in 2 days and arsenic exposure at 0.1 µM. Gene expression was calculated through mean 2−DDCt values and then normalized to the water vehicle control group and analyzed via a two-tailed t-test (*p<0:05; N= 3). Note: ATP, adenosine triphosphate; fc, fold change; LC10, lethal concentration for 10%; LC50, lethal concentration for 50%.

### Sodium arsenite inhibits cardiomyocyte differentiation and cardiac organoid formation

The 2D cardiomyocyte differentiation model mimics the prenatal cardiomyocyte formation including the mesoderm formation from ESCs and cardiac induction which leads to the cardiomyocyte differentiation. Sodium arsenite decreased cell viability during differentiation with an LC 50 of 1.3 µM while concentrations under 1 µM did not show significant cell death in the culture. Sodium arsenite exposure inhibited cardiomyocyte differentiation with a half-maximal inhibitory concentration (EC 50) of 0.05 µM (Fig 2A). At the mRNA level, 0.05 µM sodium arsenite inhibited the expression of 2D cardiomyocyte marker genes *GATA4, cTNT, alpha actinin and NKX2-5* in the differentiated cardiomyocytes (Fig 2B).

**Figure 2.**
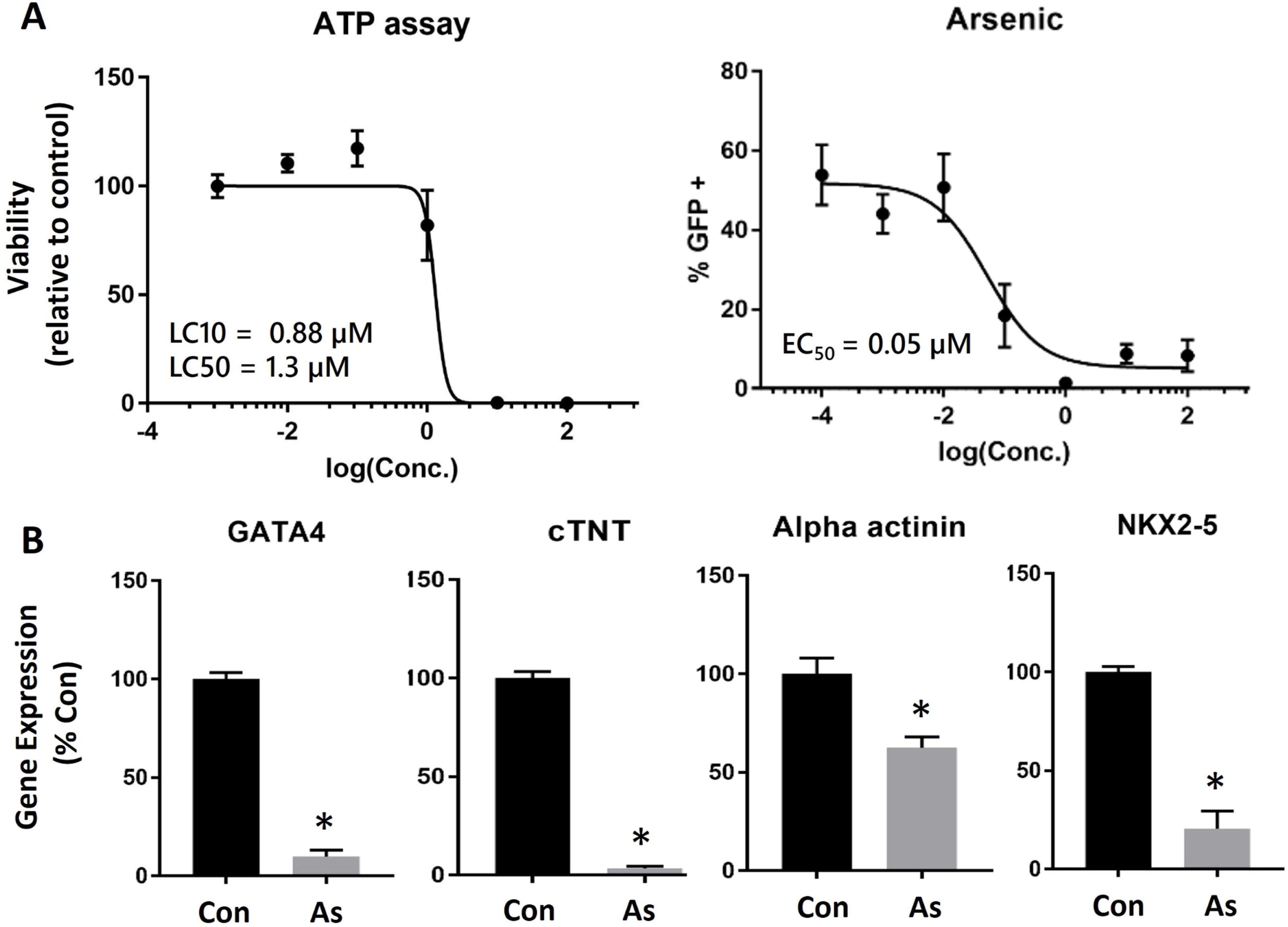
The viability of differentiating cardiomyocyte and GFP+ cardiomyocytes were assessed after 8 d of exposure to As. (A) Viability was assessed using the CellTiter-Glo assay (Promega) in 96-well plates. LC10 and LC50 were calculated under nonlinear regression curvefit and equation under log(inhibitor) vs. response in Prism software (version 10; Graphpad). (n=4–5 per group). GFP+ cardiomyocyte area was quantified by IncuCyte live cell image analysis at 8 d and EC50 was calculated under nonlinear regression curvefit and equation under log(inhibitor) vs. response in Prism software (version 10; Graphpad). (n=5 per group). (B) NKX2-5^eGFP/w^ hESCs were exposed during the time periods D0–8 and RT-PCR confirmation of the expression of selected genes associated with cardiomyocytes at 8 d of As exposure. Gene expression was normalized to the water vehicle control group and analyzed via two-tailed t-test (*p<0:05; N= 3). Note: ATP, adenosine triphosphate; Conc, concentration; EC50, median effective concentration; GFP, green fluorescent protein; LC10, lethal concentration for 10%; LC50, lethal concentration for 50%.

To further characterize the effect of sodium arsenite in the cardiac organoid model, the NKX2-5 eGFP/w hESCs were differentiated into 3D cardiac organoids and sodium arsenite was added during medium changes during the 8 days of differentiation. Expression of cardiomyocyte markers (*NKX2-5, TNNT2, TBX20, AND NFATC1*) and endothelial cell (PECAM) and fibroblast (DDR1) marker genes was higher in 3D cardiac organoids compared with expression in 2D differentiated cardiomyocytes (Fig 3A). At the mRNA level, 0.1 µM sodium arsenite inhibited the genes associated with the cardiac organoid development including cardiomyocyte markers (*NKX2-5, TNNT2, TBX20, and NFATC1*). Endothelial cell (PECAM) and fibroblast (DDR1) marker genes were also suppressed in presence of sodium arsenite exposure. Sodium arsenite exposure showed a dose-dependent organoid size decrease at 1 µM and 10 µM (Fig 3B). To determine the sodium arsenite effect on the cardiac organoid beating frequency, during cardiac organoid differentiation, sodium arsenite (0.1 µM) was added and the beat per min was not dramatically changed at the end of the 8 days cardiac organoid formation (Fig 3C).

**Figure 3.**
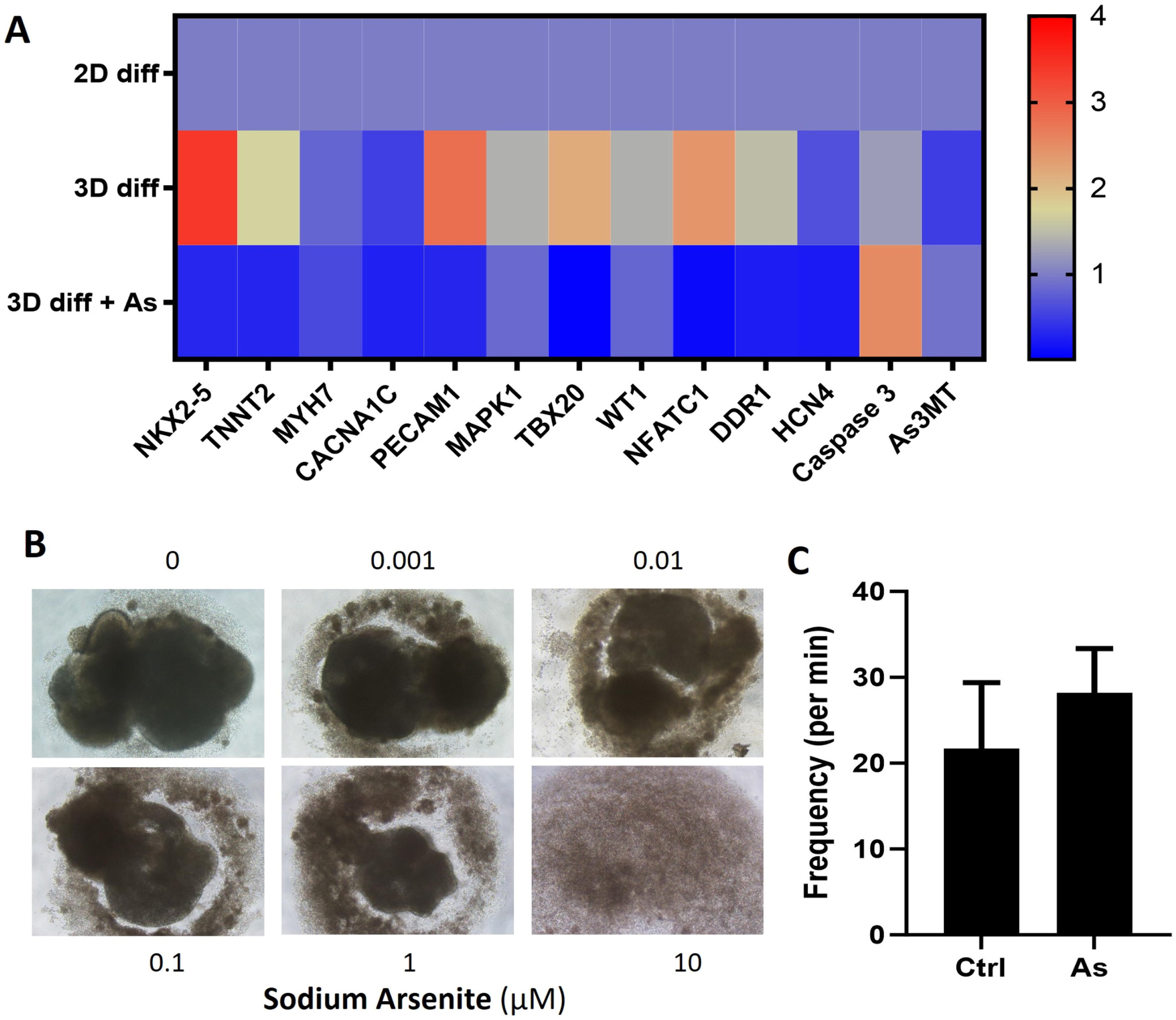
Cardiac organoid viability and gene expression in the presence of As exposure during differentiation. (A) RT-PCR confirmation of the expression of selected genes associated with cardiomyocytes, endocardial cells, and cardiac fibroblasts 8 d of cardiac organoids treated with As comparing to the 2D cardiomyocyte differentiation and 3D cardiac organoid water vehicle control group. (B) The differentiating cardiac organoids (n=3 per group) were assessed in presence of D8 exposure to As. (C) cardiac organoid beat per min was quantified using the spiky plugin in the ImageJ software with and without sodium arsenite exposure. (n=6 per group)

In addition, the effects of sodium arsenite exposure during cardiomyocyte proliferation and maturation were investigated-mimicking the perinatal stage of heart development, cells were exposed to sodium arsenite for 72 hours in an MTT assay. Sodium arsenite at concentrations of 0.001, 0.01, 0.1, and 1 µM did not significantly affect cardiomyocyte proliferation, whereas 10 µM sodium arsenite markedly inhibited proliferation (Fig 4A). In the presence of a lipid acid–rich cardiomyocyte maturation medium, 2 weeks of sodium arsenite exposure at 0.001–1 µM significantly suppressed the expression of genes involved in fatty acid oxidation, including CHKB-CPT1B and SLC25A20, as well as cardiomyocyte maturation markers α-actinin and MYH7(Fig 4B).

**Figure 4.**
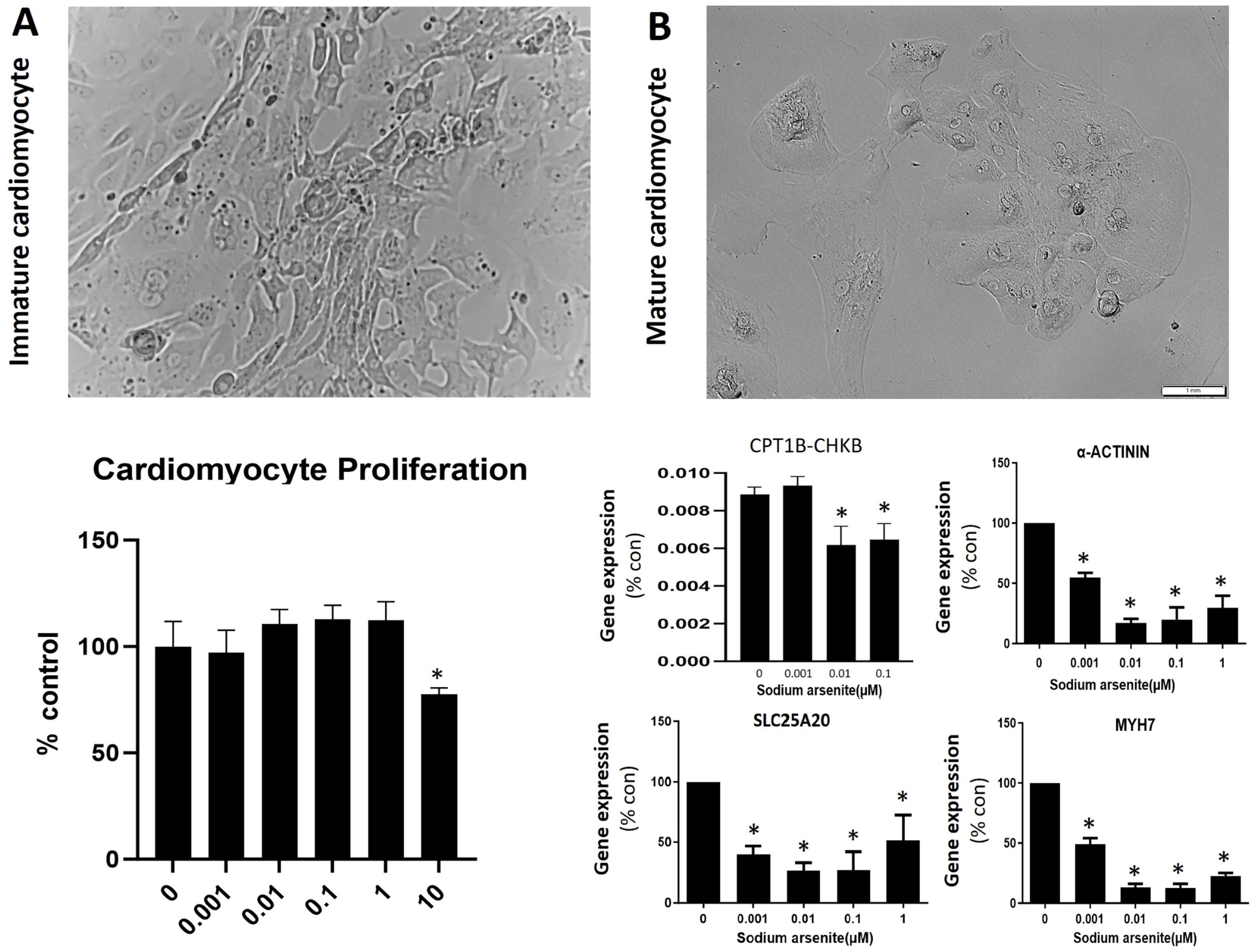
Effects of arsenic on cardiomyocyte proliferation and maturation. Day 8 cardiomyocytes were dissociated and seeded onto Matrigel-coated 96-well plates at a density of 10,000 cells per well, followed by exposure to arsenic at 0.001, 0.01, 0.1, 1, or 10 µM for 72 hours. (A) Cell viability was assessed in the MTT assay, and data were normalized to the untreated control group (mean ± SEM, p < 0.05; N = 5). In parallel, cardiomyocytes cultured in 24-well plates were maintained in lipid-rich maturation medium and exposed to arsenic at 0.001, 0.01, or 0.1 µM for two weeks. (B) Total RNA was extracted to evaluate arsenic-induced changes in gene expression. Gene expression levels were normalized to the water vehicle control group and analyzed using one-way ANOVA followed by a Tukey’s test (p < 0.05; N = 3).

### Sodium arsenite increases zebrafish heart beating rate

To study later stages of heart development after cardiomyocyte differentiation, zebrafish embryos at 24 hpf were exposed to sodium arsenite (0.1 µM) for 24 and 48 hours. After 24 hours of exposure, the untreated group heartbeat per minute was 126 ± 39 and there was a small increase in heart rate to 164 ± 73 in sodium arsenite (0.1 µM) exposure group (Fig 5A). This increase became pronounced and statistically significant after 48 hours of exposure during which untreated group zebrafish heart rate was 167 ± 46 and sodium arsenite exposure group was 207 ± 35. Quantification of gene expression after sodium arsenite (0.1 µM) exposure after 48 hours in zebrafish implicated downregulation of transcription factor gata4 and ion channel *cacna1* while transcription factors tbx1 and ion channel *kcnh2a* did not show a significant decrease in expression (Fig 5B).

**Figure 5.**
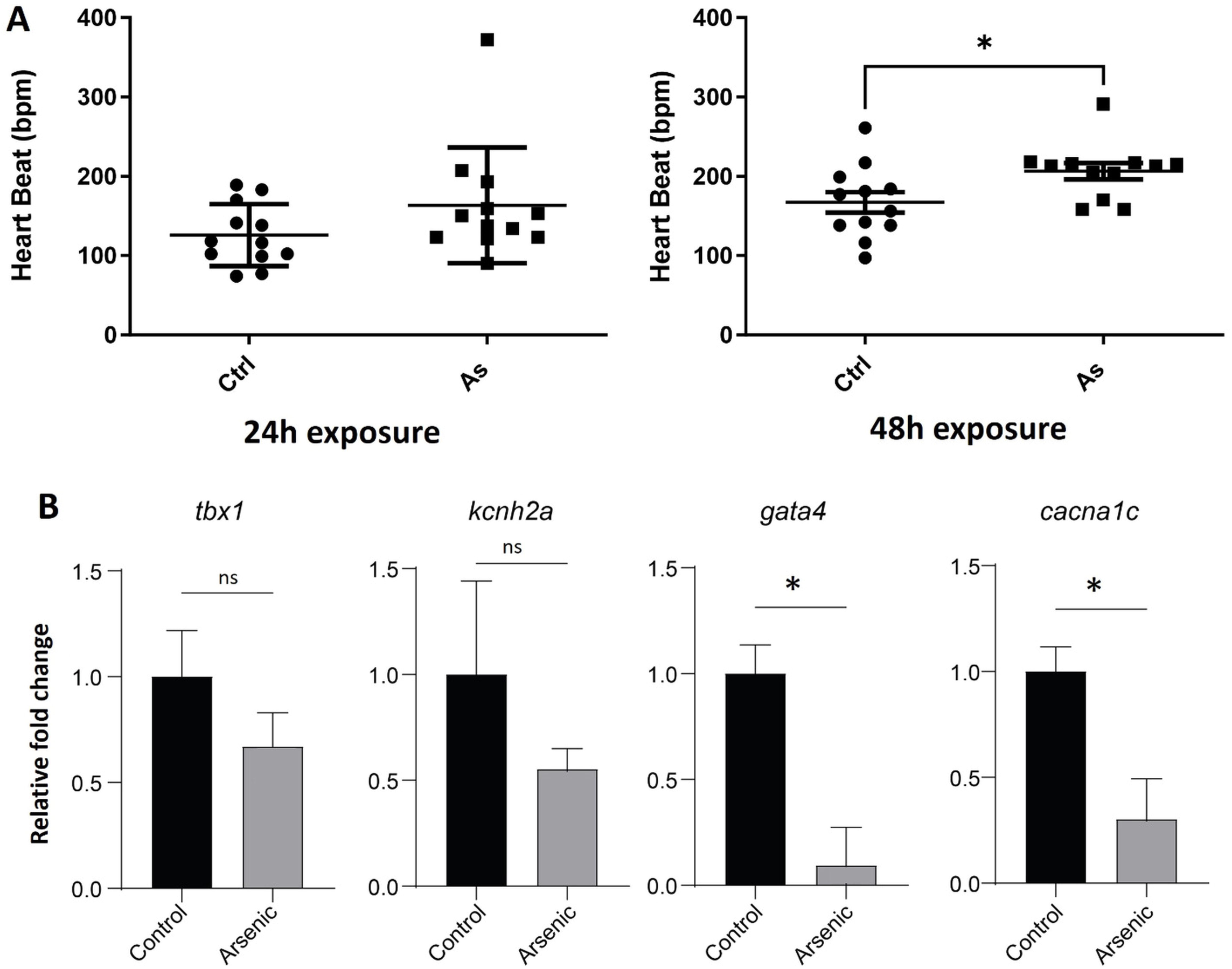
A) Effects of arsenic on heartbeat development in wildtype embryos. Embryos were sorted and placed into a 6-well plate containing 8 mL of E3 buffer, followed by exposure to 0.1 µM arsenic at 24 hpf until either 24 hour or 48 hour exposure. Data was analyzed using excel and ImageJ to determine the average heartbeat in beats per minute (bpm) of the fish. Data are the mean ± SD, with individual embryos plotted. *p < 0.05; N= 10. The 48 hpf exposure shows significance, implying arsenic effects the average heartbeat (bpm) in zebrafish when exposed for longer. B). The expression fold change of *gata4, tbx1, kcnh2a,* and *cacna1c* over age-matched untreated embryos was calculated. Mean ± SD of 3-4 technical replicates from pooled samples containing 25 different embryos are plotted. P values are from an unpaired t-test with Welch correction.

## Discussion

Disruption of cardiac development at the cell and molecular level can lead to heart development defects. To translate these insights into predictive and regulatory frameworks, it is essential to establish a testing battery that systematically covers key cellular events of heart development-including mesoderm induction, cardiac progenitor specification, differentiation, morphogenesis, and functional maturation. Such a platform would enable high-content screening of environmental toxins and pharmaceuticals for developmental cardiotoxicity, while also revealing mechanistic pathways underlying structural and functional defects. In this study, the in vitro differentiation models utilizing human embryonic stem cells have established a human-relevant system for characterizing cardiac developmental toxicology at various developmental stages. Our 3D EB screening platform illuminates the impact of chemicals on germ layer formation, while the 2D cardiomyocyte differentiation method enabled us to pinpoint the effect of sodium arsenite primarily during the cardiac induction developmental stages following germ layer formation. The in vitro cardiomyocyte proliferation and maturation assays parallel the perinatal phase of human heart development, when cardiomyocytes proliferate and begin metabolic maturation, making it highly sensitive to environmental disruptions that can cause lasting structural and functional defects. We also observed that these effects consequently resulted in heart development related gene expression change and functional alterations like heart beating rhythm in the zebrafish model. We deduced that sodium arsenite exposure 24–72 hours hpf period in zebrafish represents a critical window of heart morphogenesis, during which the primitive linear tube undergoes looping, chamber formation, and functional maturation marked by rhythmic contractions and valve development.

In the context of cardiac development, cardiac progenitor cells emerge from the lateral plate mesoderm. While high doses of sodium arsenite have been shown to disrupt mesoderm formation in iPSC differentiation models [20], the low exposure levels in the current study did not inhibit the expression of mesoderm-related genes. However, we observed that the low dose sodium arsenite exposure inhibited cardiomyocyte differentiation and cardiomyocyte maturation which are key cellular events during human heart development. Our findings parallel previous study showing that sodium arsenite disrupts cardiomyocyte differentiation in a mouse embryonic stem cell into cardiomyocytes differentiation model [21]. By systematically covering the key cellular events involved in cardiac organoid formation and cardiomyocyte maturation, our study provides a comprehensive assessment of developmental cardiotoxicity. Notably, we identify a previously unrecognized effect of sodium arsenite on fatty-acid oxidation, a metabolic process essential for the transition to mature, energy-efficient cardiomyocytes. This finding highlights an innovative mechanism whereby sodium arsenite disrupts bioenergetic metabolism in developing cardiac tissue.

Studies have indicated that acute sodium arsenite exposure in cancer treatment can result in heart function abnormalities, such as prolonged QT intervals, QRS complex broadening, T-wave flattening, arrhythmia, and atrioventricular block [22]. However, the long-term impact and consequences of low sodium arsenite exposure during development remain unclear. Our study noted that sodium arsenite exposure disrupts multiple events including cardiac induction, cardiomyocyte maturation and increased heart beating in the zebrafish model. The coordinated downregulation of NKX2-5, TNNT2, TBX20, NFATC1, PECAM1, and DDR1 indicates profound disruption of multiple essential processes in heart development. NKX2-5 and TBX20 are core cardiac transcription factors that drive early cardiomyocyte lineage specification and chamber formation[23]; their suppression reflects impaired commitment to the cardiac fate and defective morphogenesis. Reduced TNNT2 expression further suggests compromised sarcomere assembly and contractile maturation, weakening the structural and functional integrity of emerging cardiomyocytes[24]. Downregulation of NFATC1, a key regulator of endocardial cushion and valve development[25], points to impaired endocardial differentiation and intracardiac signaling. Likewise, decreased PECAM1 expression indicates disruption of endothelial development and vascular-like organization[26], both of which provide structural support and paracrine cues essential for myocardial maturation. Finally, reduced DDR1, a collagen-activated receptor involved in cell–matrix adhesion[27], suggests defective extracellular matrix remodeling and tissue organization. Together, these gene-level impairments represent a multi-layered failure of cardiac lineage specification, structural assembly, endocardial–endothelial support, and tissue organization, ultimately leading to abnormal heart development. In zebrafish, the transcription factors GATA4 showed a significant reduction in their expression levels. GATA4 functions upstream of and in cooperation with NKX2-5 as part of the highly conserved GATA-NKX axis [28]. In zebrafish, inhibition of the activity of Gata factors leads to a reduced level of *nkx2.5* and a block in cardiac progenitor cell migration. Therefore, reduced expression of *gata4* could indicate a defect in the initiation of the main cardiac regulatory network in zebrafish.

The downregulation of ion channel gene *kcnh2a* can also provide mechanistic insights into arsenic cardiotoxicity. In zebrafish, *kcnh2a* is primarily expressed in the epidermis during development and overexpression of this gene leads to long fins [29, 30]. Reduction in *kcnh2a* due to genetic mutations are associated with cardiomyocyte defects, consistent with our findings [31]. Reduction in the *kcnh2a* gene in zebrafish are typically associated with prolonged QTc interval[32]; however, our studies suggest that As increases heartrate. Given these genes are expressed in other tissues and that our gene expression was measured of the whole embryo, future studies utilizing cell-type transcriptomics or tissue-targeted delivery of As are warranted.

The U.S. EPA has set a Reference Dose (RfD) for inorganic arsenic at 0.0003 mg/kg/day, and the limit for arsenic in drinking water is 10 ppb. The association between arsenic contamination in drinking water and congenital heart disease has been documented globally [14]. Our study not only corroborates the link between arsenic exposure and early onset congenital heart disease but also provides vital data for risk assessment concerning low-dose arsenic contamination within a human-relevant model. While the cellular findings highlight the effects of arsenic on cardiomyocyte development, it is also important to consider that arsenic undergoes metabolic conversion into various metabolites, including inorganic arsenicals, monomethylated arsenicals, and demethylated arsenicals [33], each of which may exhibit distinct toxicokinetic and biological properties. Further investigation of arsenic metabolites is required to fully elucidate the mechanisms underlying developmental cardiotoxicity under physiological conditions.

In conclusion, our findings demonstrate that arsenic inhibits cardiomyocyte differentiation and maturation, mimicking the effects observed during prenatal heart development. Arsenic exposure also disrupts heart rate regulation in zebrafish embryos. The combined use of pluripotent stem cell–derived 2D assays, 3D cardiac organoids, and zebrafish embryo models provide a tiered testing strategy that recapitulates key events of cardiac development and offers valuable insights into the critical windows during which low-dose arsenic exposure impairs cardiomyocyte formation and function.

## Acknowledgements

This research was supported in part by the NIH, National Institute of Environmental Health Sciences. The contributions of the NIH author(s) are considered Works of the United States Government. The findings and conclusions presented in this paper are those of the author(s) and do not necessarily reflect the views of the NIH or the U.S. Department of Health and Human Services. This manuscript is the result of funding in whole or in part by the National Institutes of Health. It is subject to the NIH Public Access Policy. Through acceptance of this federal funding, NIH has been given a right to make this manuscript publicly available in PubMed Central upon the Official Date of Publication, as defined by NIH.

